# A method for downstream analysis of gene set enrichment results facilitates the biological interpretation of vaccine efficacy studies

**DOI:** 10.1101/043158

**Authors:** Yan Tan, Jernej Godec, Felix Wu, Pablo Tamayo, Jill P. Mesirov, W. Nicholas Haining

## Abstract

Gene set enrichment analysis (GSEA) is a widely employed method for analyzing gene expression profiles. The approach uses annotated sets of genes, identifies those that are coordinately up‐ or down-regulated in a biological comparison of interest, and thereby elucidates underlying biological processes relevant to the comparison. As the number of gene sets available in various collections for enrichment analysis has grown, the resulting lists of significant differentially regulated gene sets may also become larger, leading to the need for additional downstream analysis of GSEA results. Here we present a method that allows the rapid identification of a small number of co-regulated groups of genes – “leading edge metagenes” (LEMs) - from high scoring sets in GSEA results. LEM are sub-signatures which are common to multiple gene sets and that “explain” their enrichment specific to the experimental dataset of interest. We show that LEMs contain more refined lists of context-dependent and biologically meaningful genes than the parental gene sets. LEM analysis of the human vaccine response using a large database of immune signatures identified core biological processes induced by five different vaccines in datasets from human peripheral blood mononuclear cells (PBMC). Further study of these biological processes over time following vaccination showed that at day 3 post-vaccination, vaccines derived from viruses or viral subunits exhibit patterns of biological processes that are distinct from protein conjugate vaccines; however, by day 7 these differences were less pronounced. This suggests that the immune response to diverse vaccines eventually converge to a common transcriptional response. LEM analysis can significantly reduce the dimensionality of enriched gene sets, improve the identification of core biological processes active in a comparison of interest, and simplify the biological interpretation of GSEA results.

**Author Summary:** Genome-wide expression profiling is a widely used tool to identify biological mechanisms in a comparison of interest. One analytic method, Gene set enrichment analysis (GSEA) uses annotated sets of genes and identifies those that are coordinately up‐ or down-regulated in a biological comparison of interest. This approach capitalizes on the fact that alternations in biological processes often cause the coordinated change of a large number of genes. However, as the number of gene sets available in various collections for enrichment analysis has grown, the resulting lists of significant differentially regulated gene sets may also become larger, leading to the need for additional downstream analysis of GSEA results. Here we present a method that allows the identification of a small number of co-regulated groups of genes – “leading edge metagenes” (LEMs) – from high scoring sets in GSEA results. We show that LEMs contain more refined lists of context-dependent biologically meaningful genes than the parental gene sets and demonstrate the utility of this approach in analyzing the transcriptional response to vaccination. LEM analysis can significantly reduce the dimensionality of enriched gene sets, improve the identification of core biological processes active in a comparison of interest, and facilitate the biological interpretation of GSEA results.

## Introduction

Changes in the state of a cell or tissue are often reflected by the alteration of large numbers of genes. Analysis of large-scale datasets from yeast [1] to humans [2] demonstrates that changes in gene expression accompanying a biological shift in the cell state are organized into “modules” or sets of genes that play a functional role in executing a particular biological process. However, the change in gene expression in many transcripts contained in such modules can be of such small magnitude that they would be hard to detect over experimental noise when considered individually [3,4]. As a result, several computational tools have been developed to detect the coordinate up-regulation of a program of genes [5] with robust statistical measures of significance, even though the absolute change in expression of any constituent gene in the set of genes may be small.

One widely used approach is gene set enrichment analysis (GSEA) [6], which tests whether a set of genes of interest are randomly distributed throughout a rank-ordered list of genes (usually generated by comparison of the gene expression profiles of two phenotypic classes) or are over-represented at the top or bottom of the list. The latter finding then leads to the inference that this set of genes is related to the underlying biology of the two phenotypes in question. The power of GSEA and other analytic enrichment tools to yield insights into biology is critically dependent on the number and quality of databases of sets of genes, which are tested for enrichment in the phenotypic comparison of interest. Some databases, like Gene Ontology [7] or TRANSFAC [8], represent collections of genes generated *a priori*, without reference to specific experiments. Others, such as MSigDB [9] contribute a large number of gene sets curated from experimentally derived expression profiles corresponding to cell states and biological perturbations.

However, the increasing numbers of gene sets that become available for analysis can lead to the need for additional downstream analysis of GSEA results. The first challenge arises when a list of enriched gene sets contains considerable redundancy, i.e., multiple sets with a subset of genes – “sub-signatures” – in common. For example, results of GSEA may contain multiple gene sets each of which contains a signature of genes corresponding to a common biological process such as proliferation or interferon response, even though the experiments that elicited proliferation or interferon response were quite distinct. While the annotation of the gene sets themselves can provide considerable biological insights, it may not be immediately apparent whether the enrichment of two or more gene sets is due to the presence of the same set of sub-signatures in multiple gene sets that “explains” their enrichment.

A second challenge is that as increasing numbers of gene sets become available for use with GSEA, the results of an analysis can sometimes include tens or hundreds of significantly enriched gene sets requiring considerable investigator review. The interpretation of GSEA results would therefore be facilitated by the development of tools to reduce the dimensionality of GSEA results. One way to do this would be to identify coordinately up-regulated sub-signatures contained within several enriched gene sets corresponding to the biological themes present in the phenotypic comparison of interest.

To meet this need, we have developed an analysis approach to identify sub-signatures of genes which we term “leading edge metagenes” (LEMs) that are both common to multiple, significantly enriched gene sets and coordinately enriched in a phenotypic comparison of interest. The “leading edge” of a gene set consists of those genes that “drive” the enrichment score in a GSEA analysis, and represent a rich source of biologically important genes (see Methods). We show that the LEMs are more significantly enriched for biologically related genes than the parent gene sets or their leading edges. We apply this approach to the analysis of the transcriptional response induced by five different vaccines measured in peripheral blood mononuclear cells (PBMC). By examining the results of a GSEA analysis in the space of the LEMs, we can more clearly see that while different vaccines initially elicit distinct, biologically coherent patterns of gene expression three days after vaccination, these transcriptional patterns of response become more similar over time.

## Results

### Overview of the leading edge metagene method

We developed an approach to identify groups of genes – termed “leading edge metagenes” (LEMs) – that are both associated with a phenotype of interest and shared between multiple gene sets enriched in that phenotypic comparison. We reasoned that groups of genes that are co-regulated in the phenotype of interest and also present in multiple gene sets are likely to represent the core sub-signatures of genes related to distinct biological processes or pathways. Our approach leverages the notion of the “leading edge genes” in a GSEA analysis [6], which are the genes whose expression profiles are most highly correlated with the phenotype distinction in a comparison of biological states and thus drive the GSEA enrichment statistic. We present here an overview of the leading edge metagene (LEM) method that is summarized in Fig. 1, and give more details in Methods. The LEM method and source code are available on GitHub (https://github.com/lamarck2008/LEM).

**Step 1:** Use GSEA to identify the enriched gene sets in a phenotypic comparison of interest, such as gene expression profiles of PBMC samples before and after vaccination. Extract the leading edge genes from each enriched gene set.
**Step 2:** Construct a sparse *n* by *m* matrix *M* of genes by gene sets. The entries in each column give the corresponding leading edge gene’s correlation with the phenotype distinction in the data set. Note that if a gene is not in the gene set’s leading edge its entry will be 0.
**Step 3:** Apply nonnegative matrix factorization (NMF) [10,11] to *M*. This will yield a product of two matrices *W* × *H* that approximates *M*, where the entries of the columns of *W* indicate the contribution of each gene to the corresponding metagene and the *H* matrix represents the gene sets in the space of metagenes.
**Step 4:** Filter the metagenes, (columns of the matrix *W*), i.e., set to 0 the entries with a coefficient that falls below a threshold. As each metagene is a positive linear combination of all the genes, a small coefficient indicates negligible contribution to the metagene. This has the effect of removing the corresponding genes from that metagene.
**Step 5:** Define the *leading edge metagenes* (LEMs) for each column in the *W* matrix by assigning each gene with non-zero entries in its row to the leading edge metagene with the largest coefficient.

**Figure 1.**
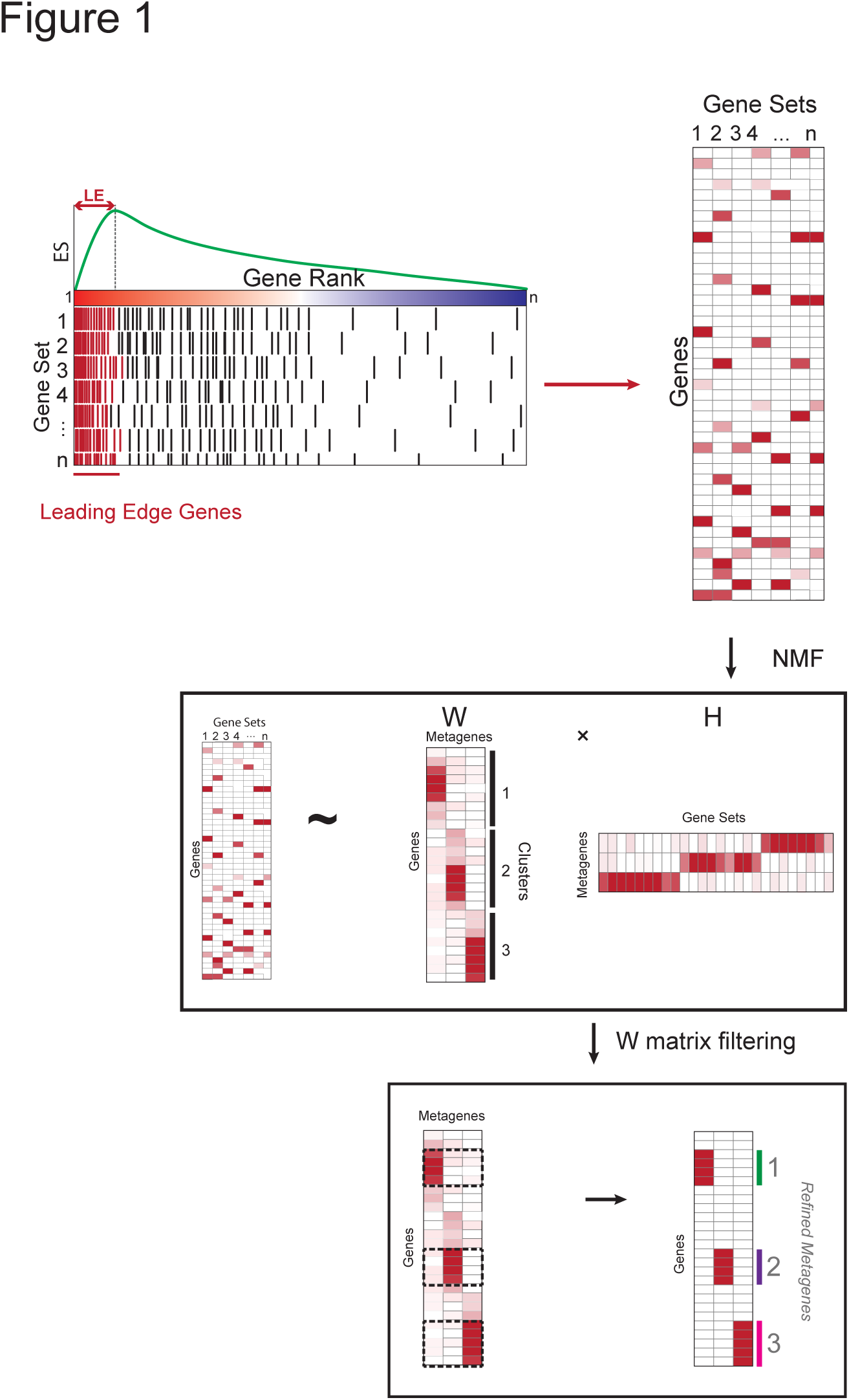
Schema of leading edge metagene analysis.

### Gene set enrichment analysis of the transcriptional response to YFV vaccination

We applied this approach to the gene set enrichment analysis of the transcriptional response to vaccination. Transcriptional profiling of the vaccine response has been used to identify biological processes associated with different vaccines and to develop predictors of protective immunity following vaccination [12–15]. The changes in gene expression in PBMC following yellow fever vaccine (YFV) vaccination of healthy volunteers have been well-studied [12,14], and provided a useful test case in which to apply LEM analysis. We studied a dataset of gene expression profiles of PBMC from healthy volunteers (*n* = 15) either before (day 0) or after (day 7) vaccination with YFV-17D, a live attenuated viral vaccine. We ranked genes by their differential expression in day 7 vs. day 0 and performed GSEA using the Immune Signatures C7 collection of MSigDB (ImmuneSigDB), a compendium of almost 2,000 signatures [16] curated from experimentally derived gene expression profiles in the immunology literature. We identified 481 gene sets that were significantly enriched at day 7 following YFV vaccination relative to day 0 (Fig. 2 A). Many of the enriched gene sets were those that would be expected to correlate with vaccination-induced changes in gene expression, such as those related to inflammation, cell proliferation, and response to virus.

**Figure 2.**
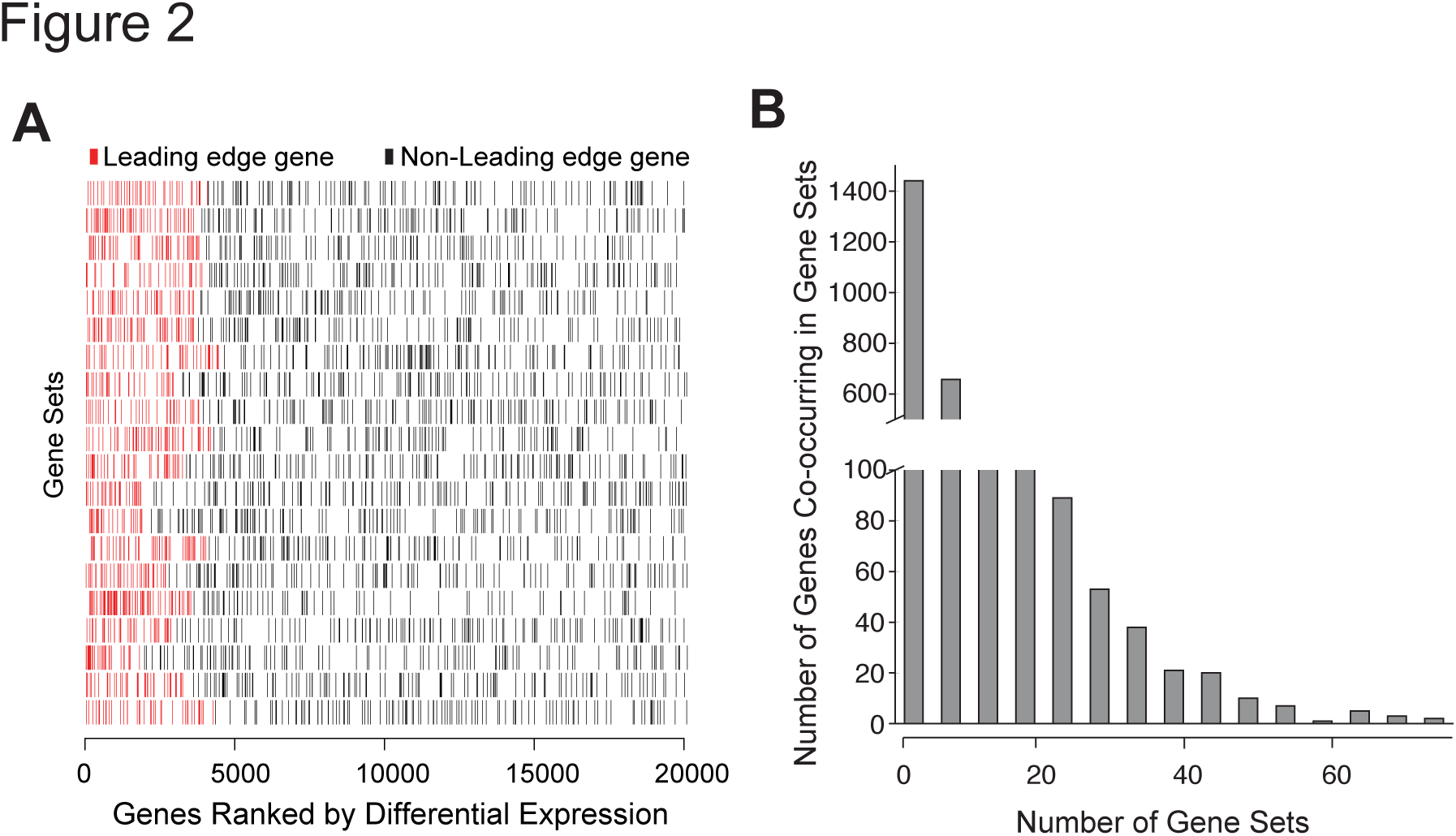
GSEA of the transcription response to YFV vaccination identifies leading edge genes. **(A)** GSEA of genes up-regulated in PBMC after YFV vaccination in healthy volunteers using the MSigDB C7 Immune Signatures Database. Each gene set in the top 20 of gene sets are shown as rows of “barcodes” representing the position of each gene the gene set on the ranked list of genes differentially expressed post‐ vs. pre-vaccine (X-axis). Leading edge genes (red) are those that occur before the maximal enrichment score for each gene set. **(B)** Frequency histogram of the number of leading edge genes co-occurring in multiple gene sets.

We extracted the leading edge genes from all 481 enriched gene sets and assessed the frequency of co-occurrence of genes in gene sets (Fig. 2 B). Of the 2821 leading edge genes present in any of the enriched gene sets, the vast majority (75%) were present in 10 or fewer gene sets, and the resulting matrix of gene sets and leading edge genes was sparse with 98% of the entries equal to 0. (S1 Fig.)

### Identifying LEMs in the transcriptional response to YFV vaccination

We applied an NMF consensus clustering method to estimate the appropriate number of metagenes in the leading edge sparse matrix [11]. As is suggested by the consensus matrix in Fig. 3 A, there are three gene clusters represented in the leading edge sparse matrix suggesting that a rank 3 *W* matrix is the best low dimensional approximation. A simple inspection of coefficients in each of the columns of the *W* matrix suggests that most genes have very small coefficients and only a small fraction of genes have significantly large coefficients (Fig. 3 B). To filter out genes with negligible contributions to the metagenes, we fit three exponential distributions to the coefficients of each metagene in *W*. As is shown in Fig. 3 B, genes with coefficients below the cutoff of 1 are colored white and genes with coefficients above the cutoff are masked with a color gradient where more red indicates higher coefficient values. Genes that fall in the white regions were then filtered from the *W* matrix and the remaining genes were assigned to one of the three LEMs based as described in Methods.

**Figure 3.**
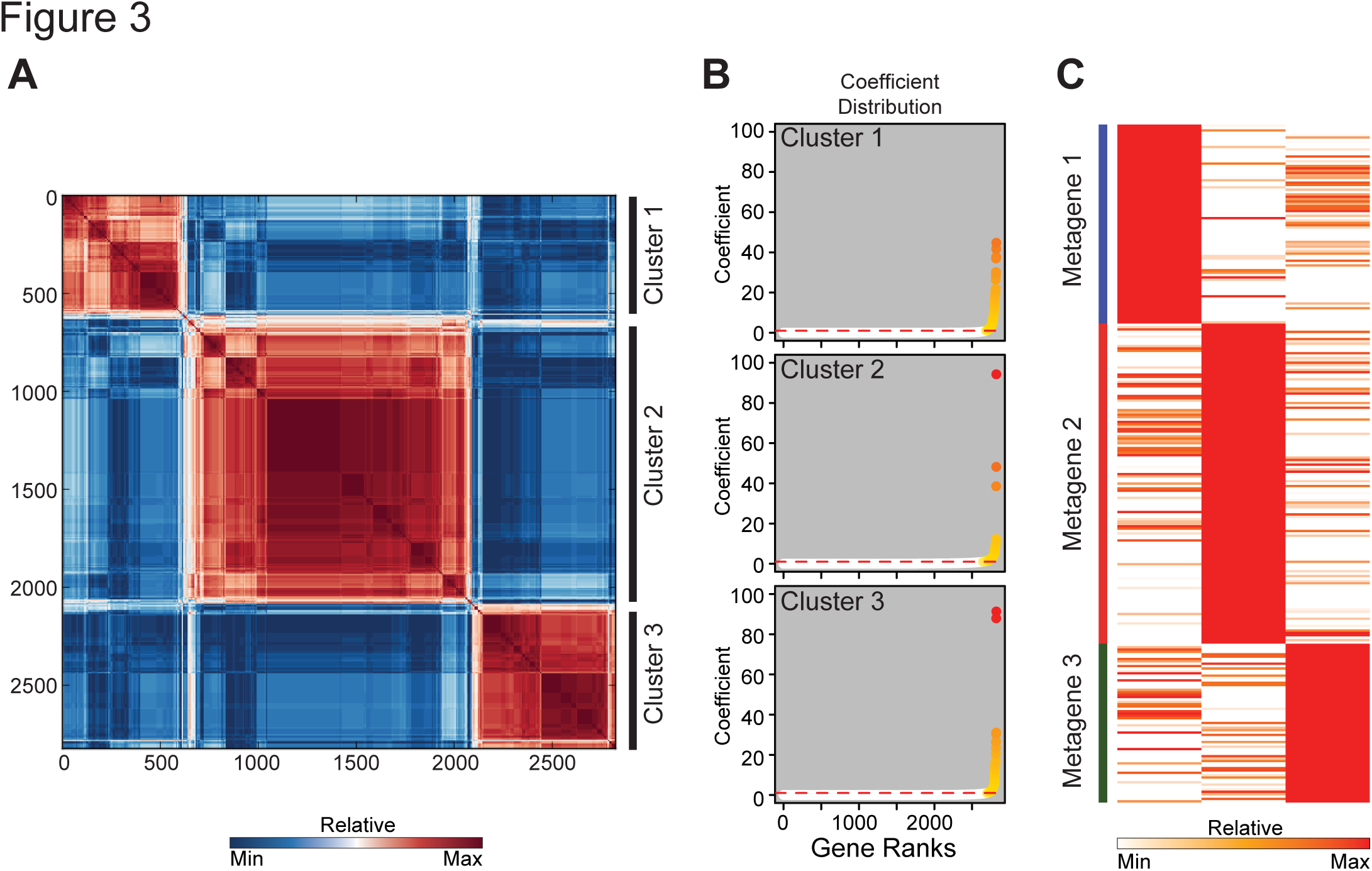
LEM analysis identifies three leading edge metagenes in the transcriptional response to YFV vaccination. **(A)** Consensus cluster matrix for gene sets membership values averaged from 100 matrices using 2821 leading edge genes from 482 gene sets enriched in day 7 post-YFV profiles compared to pre-vaccine profiles computed at *k* = 3. Red values are highly correlated; blue uncorrelated. **(B)** Coefficients of the contribution of each leading edge gene to the three clusters of genes identified in (A). Dotted red line shows the threshold of 1 which is 95 quantile based on a fitted exponential distribution. **(C)** Heatmap of the coefficients (post vs. pre vaccine) of genes assigned to each of three metagenes present in the YFV vaccine signature. Representative genes indicated on the right.

### LEMs are highly enriched for genes associated with biological processes

We reasoned that genes contained in LEMs would be a more refined list of biologically related genes than those in the gene sets from which they were extracted. Experimentally derived gene sets are generated from the comparison of two phenotypic classes, and as such are likely to entrain genes related to multiple, different biological processes as well as a variable degree of experimental noise. In contrast, LEMs, are “filtered” by virtue of appearing at the leading edge of enrichment in a phenotype of interest (here, day 7 post-vaccination) across multiple experimentally-derived gene sets. We therefore tested whether LEMs were more highly enriched for genes related to biological processes (as annotated by Gene Ontology) than their parental gene sets (Fig. 4). We tested the set of 3 LEMs for overlap with the collection of GO annotated gene lists, and determined the significance of each GO term’s overlap. We compared the P-values generated by GO term overlap with LEM genes, an equivalent number of genes randomly sampled from the original pool of leading edge genes, and genes sampled from the entire transcriptome. We found that the significance of GO term overlap was much higher in the LEMs than in the original leading edge genes or in a random set of genes. GO terms that were enriched included many with known roles in the vaccine response to attenuated viral vaccines including those related to virus response, cytokine production and proliferation. We note that these enriched GO terms are not directly interchangeable with the LEMs themselves. While the overlaps between metagenes are significant, GO term annotated genes account for only a small fraction of all the genes in each LEM (Fig. 5). LEM analysis therefore provides an effective approach to refine GSEA results to a subset of genes most related to defined biological processes.

**Figure 4.**
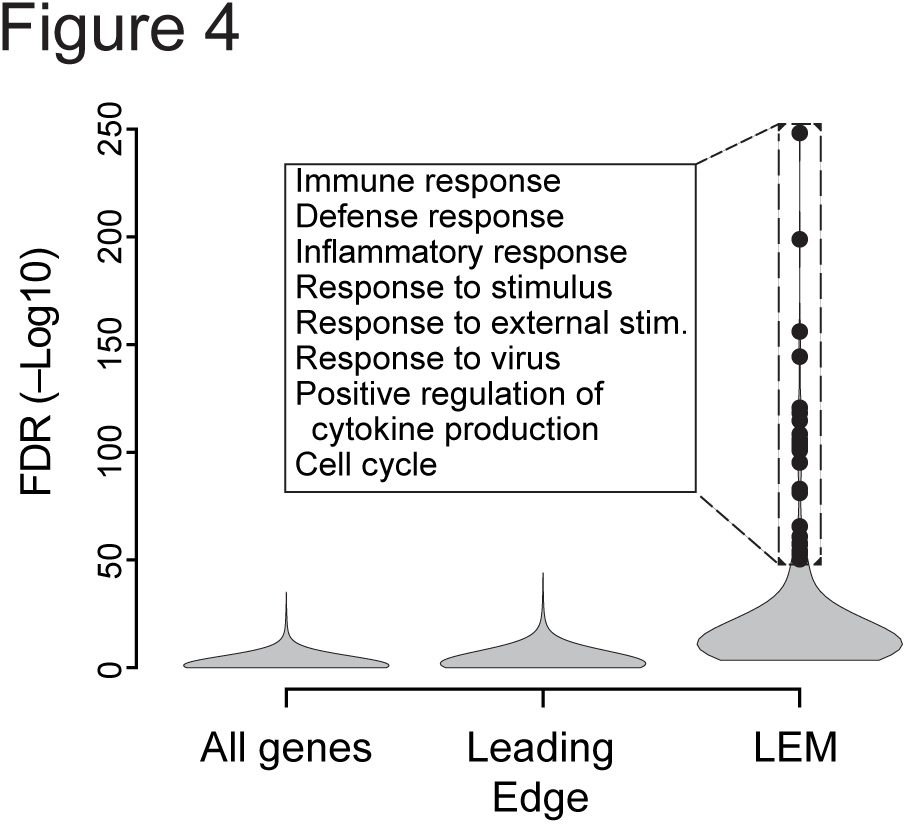
Leading edge metagenes are more enriched for biological signatures than the unrefined set of leading edge genes. Violin plots of the distribution of ‐log10 hypergeometric P values calculated from overlap of GO term-annotated genes with LEMs (right), randomly selected groups of genes of similar size from the leading edges (middle) of enriched gene sets, or the whole genome (left). Black dots represent the ‐10 * log_10_ *P* value for GO terms that enrich with a *P* value < 1e-5.

**Figure 5.**
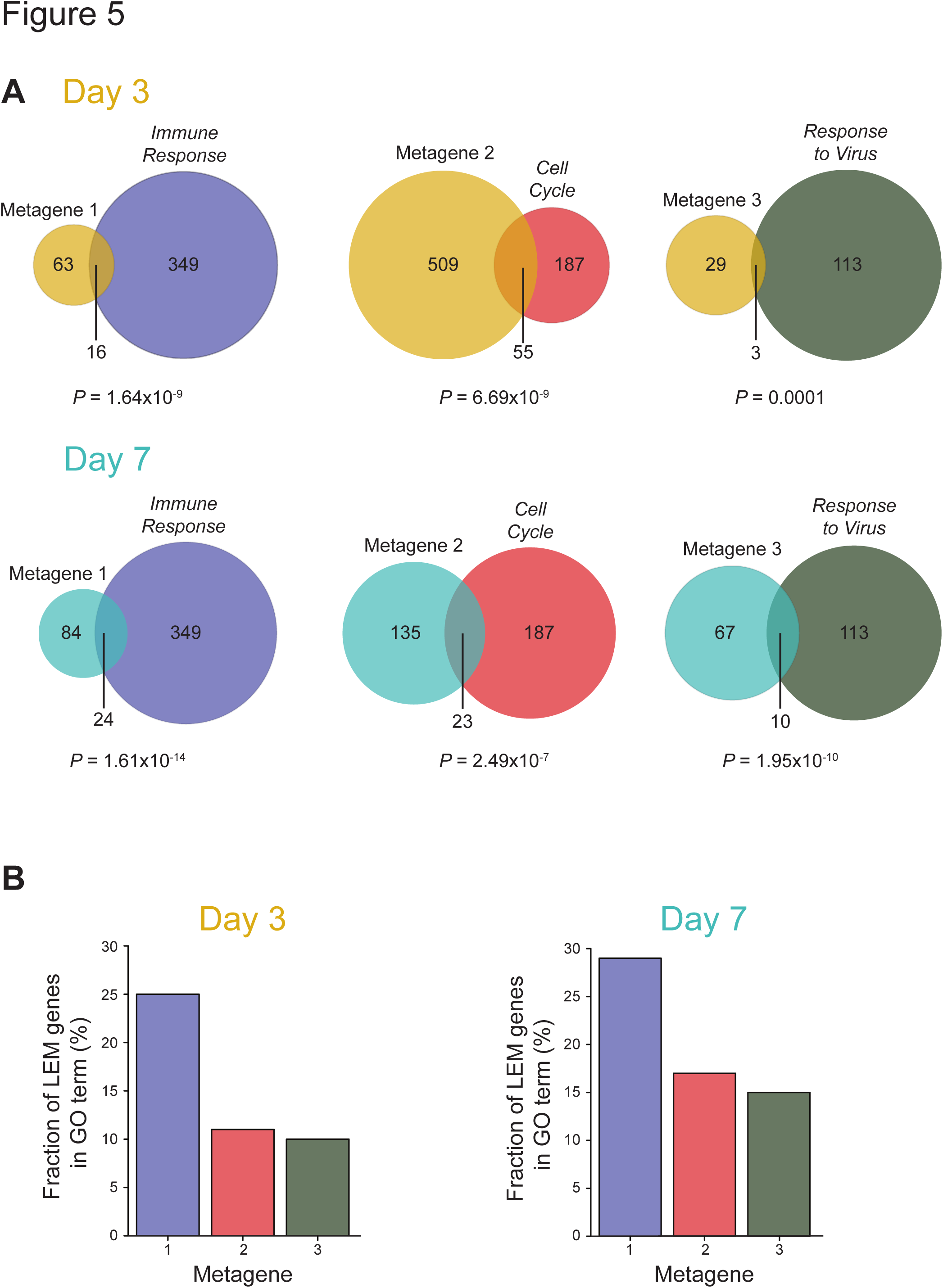
GO annotations of leading edge metagenes are not interchangeable. **(A)** Venn diagrams of gene overlaps between YFV LEMs and significantly enriched Gene Ontology terms – Immune Response (blue), Cell Cycle (red), and Response to Virus (green) – at day 3 and day 7 post-vaccination. Circles are drawn in relative proportion to the number of genes in the metagene or term. **(B)** Bar plots indicating the fraction of YFV LEM genes annotated by a particular GO term at day 3 and day 7 post-vaccination. Bar colors correspond to the colors of GO term circles in the Venn diagrams above.

### LEM analysis identifies conservation and divergence of biologic kinetics in human vaccines

We extended LEM analysis to an additional four previously published datasets of gene expression profiles from PBMC following vaccination [14]. The vaccines studied differed in immunogens and adjuvant, mechanism of action, and targeted disease. They included both polysaccharide and conjugate vaccines targeting meningococcal disease (MPSV4 and MCV4, respectively), trivalent influenza vaccine (TIV) and live attenuated virus vaccines targeting influenza (LAIV).

For each vaccine, we compared the transcriptional response at day 3 or at day 7 post-vaccination to the baseline time point at day 0 prior to vaccination for a total of 10 comparisons (2 time points × 5 vaccines). We identified significantly enriched gene sets in the post-vaccination samples again using the ImmuneSigDB [16] and extracted LEMs associated with the vaccine response for each vaccine at both time points. Initial GSEA results returned between 0 and 550 significantly gene sets that were enriched in each of the post-vaccination datasets relative to pre-vaccination samples. In nine of our ten comparisons, LEM analysis reduced these large collections of gene sets to three LEMs; analysis of the MCV4 vaccine response at day 0 versus day 7 post vaccination yielded four LEMs. We annotated this set of thirty-one LEMs using Gene Ontology (GO) terms based on the strongest overlap of a LEM with one of the ten representative GO terms (Fig. 6) identified by REVIGO [7,17]. We identified LEMs related to biological processes that have previously been shown to be involved in the PBMC transcriptional response to vaccination. For example, we found that LEMs comprised of genes related to cell cycle, immune response, and response to virus and stress were induced by several vaccines and at different time points. This is consistent with previous reports that found similar biology to be important in vaccine-induced responses [12,13,15,18,19].

**Figure 6.**
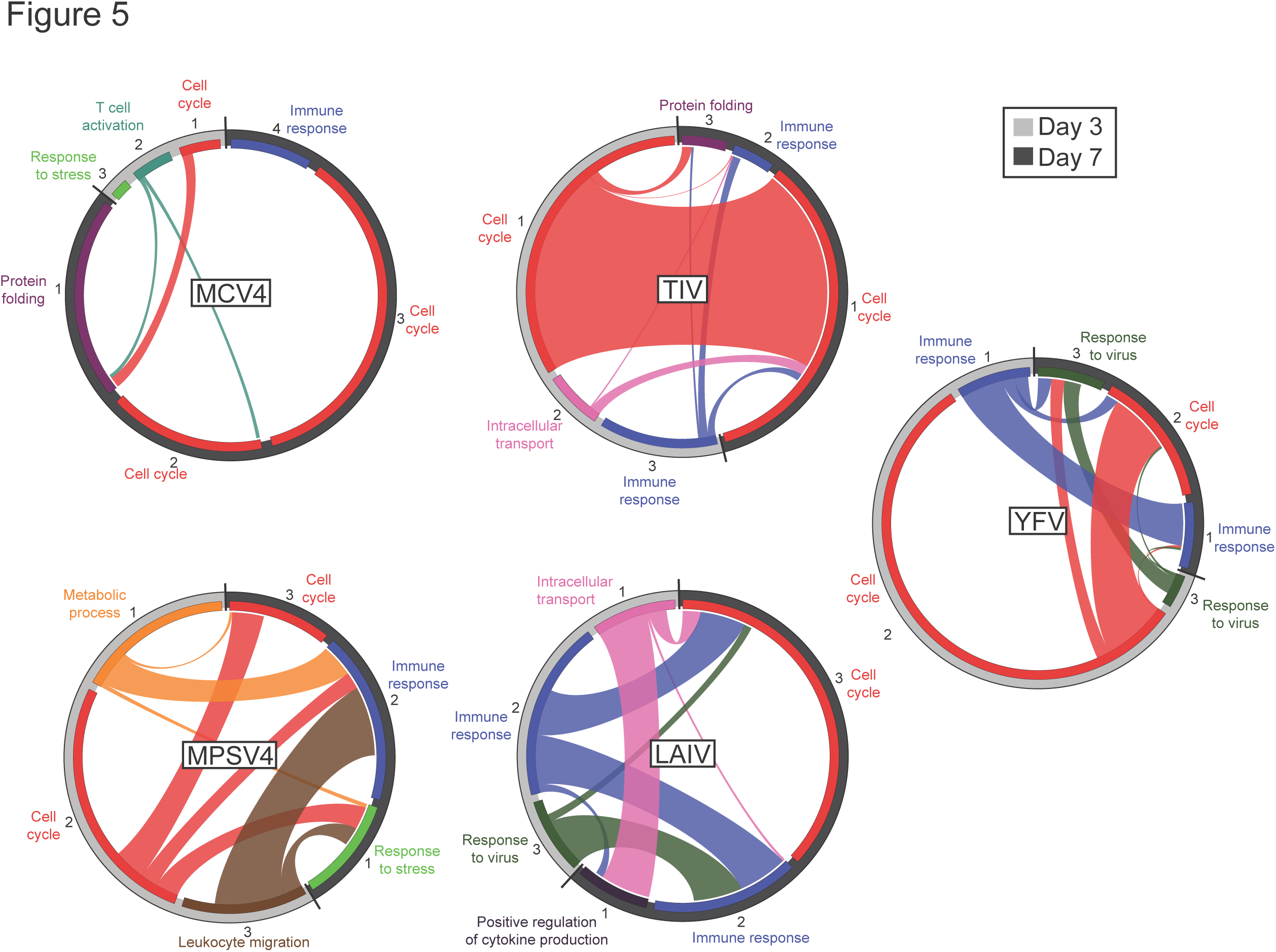
Comparison of leading edge metagenes elicited three and seven days after vaccination by five vaccines. Circos plots indicating overlap in gene membership of metagenes elicited by five vaccines (indicated in the box at the center of the plot) on day 3 (grey band) vs. day 7 (black band) post-vaccination. Breadth of the connecting ribbon is proportional to the fraction of genes shared between metagenes. Predominant biological process present in each arbitrarily-numbered metagene indicated in text beside each segment.

We next compared the LEMs that were identified at day 3 post vaccination to those found at day 7 for each vaccine by comparing the fraction of shared genes in each. We found that the degree of overlap of LEMs at day 3 and day 7 varied widely in different vaccines (Fig. 6). For example, the transcriptional response to MCV4 at day 3 induced LEMs that had minimal overlap with those for day 7. However, the closely related vaccine, MPSV4 showed a striking overlap in the LEMs between day 3 and day 7. In the case of TIV, a LEM representing cell cycle progression evident at day 3 persisted into day 7, whereas LEMs related to intracellular transport and immune response observed at day 3 showed minimal overlap with LEMs found at day 7 in the same vaccine. This suggests that while proliferation is a prominent part of the transcriptional response to TIV at day 3 and day 7, other transcriptional features shifted in this time frame. This contrasts with the LEMs elicited by vaccination by LAIV, which showed no evidence of proliferation until day 7. This is potentially related to the fact that TIV is administered intramuscularly and thus may elicit a more rapid response when measured in the blood than LAIV, which is given by the intranasal route.

### Diverse vaccines elicit distinct transcriptional responses at day 3 but become more similar at day 7

The vaccines examined vary in their composition and include live attenuated viral vaccines (YFV and LAIV), inactivated virus (TIV), carbohydrate (MPSV4), and polysaccharide-conjugate (MCV4). We therefore examined the shared and unique LEMs induced by different vaccines at day 3 and at day 7 post vaccination. We computed the significance of each pairwise overlap of LEM membership using a hypergeometric test, and visualized these comparisons as a heatmap (Fig. 7 A).

**Figure 7.**
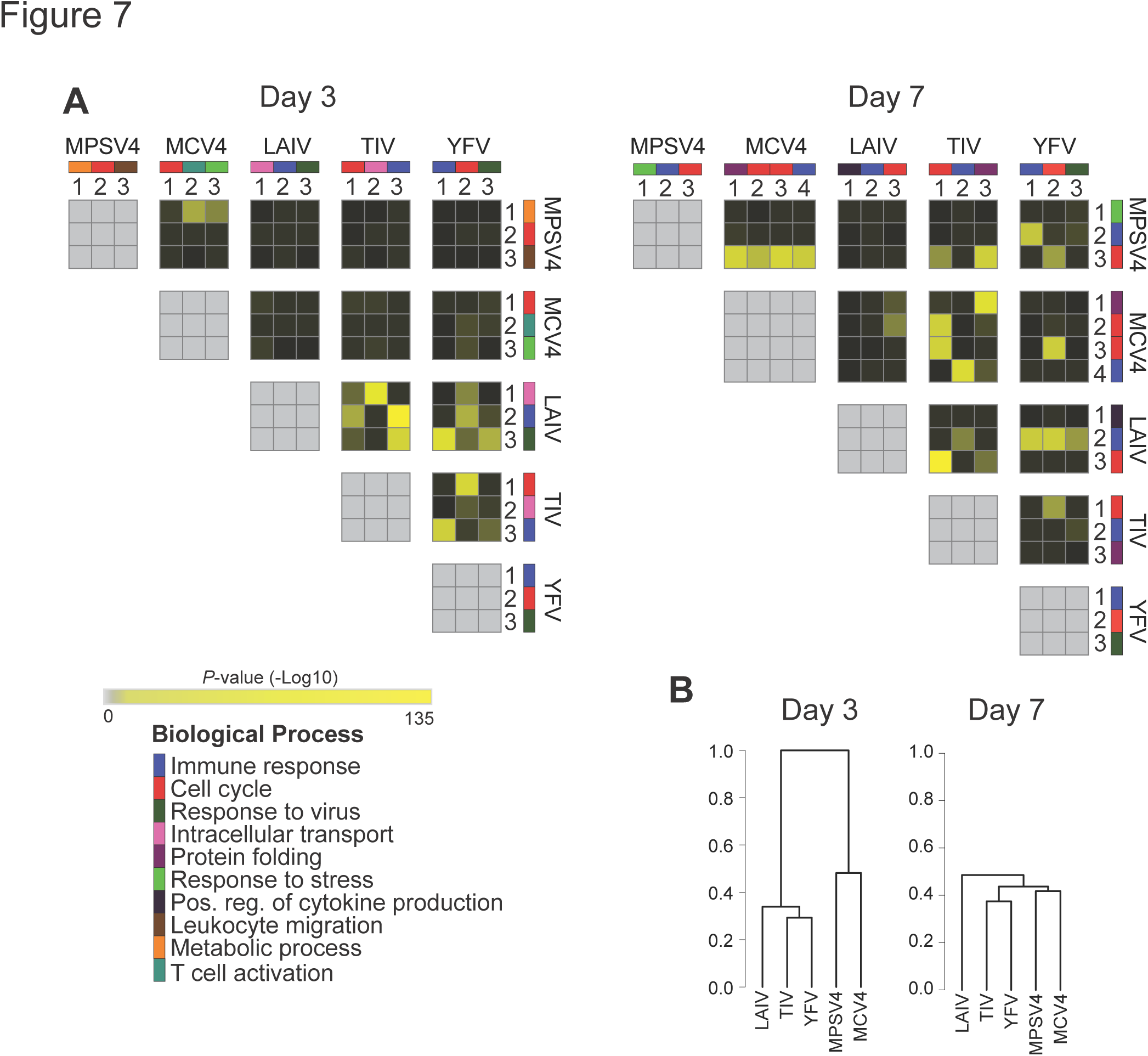
The transcriptional response in five different vaccines becomes more similar over time. **(A)** Pairwise overlap of metagenes elicited by five vaccines on day 3 post vaccine (left) or on day 7 post vaccine (right). Heatmap values correspond to the significance of the overlap (shown as ‐log10 hypergeometric *P* values). Each metagene is given an arbitrary number, and the predominant biological process present in each metagene indicated by the color key. **(B)** Hierarchical clustering of vaccines calculated by similarities in the significance of metagene overlap at day 3 post vaccine (left) or on day 7 post vaccine (right).

At day 3, we found that the LEMs elicited by a meningococcal carbohydrate vaccine (MPSV4) and a meningococcal polysaccharide conjugate vaccine (MCV4), showed significant overlap of LEMs related to T cell activation, stress response and metabolism. Surprisingly although both meningococcal vaccines elicited LEMs containing genes related to proliferation, the sets of genes comprising the proliferation LEMs themselves showed minimal overlap, possibly suggesting different mechanisms or kinetics of cell cycle progression induced by these different vaccines. In contrast, neither meningococcal vaccine showed significant overlap with any of the LEMs in the constituent genes elicited by viral vaccines. However, there was significant overlap in the LEMs elicited by the inactivated and live attenuate viral vaccines including LEMs related to cell cycle, intracellular transport, and the immune response. We visualized the relative distance between two vaccines using hierarchical clustering based on a distance derived from the P-values calculated by all pairwise overlap comparisons of LEMs from each of the two vaccines (see Methods). This analysis demonstrated that the meningococcal vaccines clustered in a distinct branch of the dendrogram separate from the viral vaccines (Fig. 7 B, left).

At day 7, however, we observed a much broader degree of overlap in the constituent genes between metagenes induced by different types of vaccines. In contrast to day 3, we found that multiple LEMs elicited by meningococcal protein vaccines showed similarity to those elicited by viral vaccines. For instance, the proliferation LEM induced by YFV (live attenuated vaccine) showed significant similarity to the proliferation LEM elicited by MPSV4 (*P*=8.65×10^−6^), MCV4 (*P*=6.65×10^−8^), as well as to LAIV (*P*=2.2×10−^5^) and TIV (*P*=9.16×10^−6^). Similarly an immune response and protein-folding LEM elicited by MCV4 showed significant overlap with LEMs elicited by TIV and YFV. As a result, hierarchical clustering showed a much closer degree of relatedness between all five vaccines at day 7 than was seen at day 3 (Fig. 7 B, right). Thus the patterns of LEM expression elicited by carbohydrate and protein conjugate vaccines are distinct from those elicited by viral vaccines at day 3, but become more similar at day 7.

## Discussion

Identifying the coordinate up‐ or down-regulation of biologically meaningful sets of genes has become an important tool for the analysis of gene expression data. Several large collections of biologically informative gene sets enable comprehensive annotation of experimental datasets using gene set enrichment analysis (GSEA). However, the results of GSEA may include large numbers of gene sets that are enriched in the experimental dataset and the manual biological interpretation of every one of them can prove challenging and time consuming. Our method allows the rapid identification of a small number of LEMs common to multiple gene sets and coordinately regulated in the experimental dataset of interest. This approach complements standard GSEA by identifying LEMs that “explain” the enrichment of multiple gene sets. It starts with a large group of enriched gene sets and reduces them to a smaller number of LEMs that correspond to specific biological themes present in the phenotype of interest and that are specific to the data set studied. We applied this approach to identify the biological processes that are elicited by five different vaccines, and identified both shared and vaccine-specific components of the immune response to vaccination at different time points.

Many gene sets in MSigDB are curated from expression profiles derived from experimental comparisons. As such, they often represent complex biological events, such as signatures induced by genetic perturbation or based on comparison of distinct differentiation states. It is therefore likely that many of these gene sets include multiple sub-signatures, each representing distinct biological processes. For instance, during the differentiation of effector CD8^+^ T cells from their naive precursors, there is marked up-regulation of sets of genes related to proliferation and also other genes related to effector T cell function. The GSE9650_NAIVE_VS_EFF_CD8_TCELL_DN gene set in the ImmmuneSigDB collection within MSigDB, therefore, contains genes related to mitosis (e.g., *CDK1* and *CDC34*) as well as effector genes (e.g., *GZMB* and *IFNG*). The former class of genes is likely to be shared with activated and proliferating B cells, but not the latter. This heterogeneity can present a challenge to interpretation of multiple high scoring gene sets. The approach that we have developed extracts core sub-signatures, or leading edge metagenes (LEMs), relevant to the context of the investigator’s data set. This approach achieves two goals: first, it reduces the complexity of the results of GSEA by reducing the number of gene sets that need to be evaluated, and second, it creates a refined set of LEMs in which key biological themes present in a phenotypic comparison of interest can be readily identified.

The problem of finding intersecting subgroups of genes in the leading edge of multiple gene sets is challenging because the overlaps consist of a relatively small number of genes (Fig. 2 B), resulting in very sparse gene by gene set matrices that require proper analysis to extract the relevant gene subgroups. This type of analysis can be done, for example, using matrix decomposition techniques with the important requirement that the decomposition produces binary sparse matrices so that the “components” representing the most important overlaps can in turn be interpreted as sub-signatures. Traditionally, methods such as singular value decomposition (SVD) [20] and PCA [21] have been used to decompose input matrices into relevant components that represent the most salient relationship between variables and data points. However, these methods do not produce sparse representations and are therefore less suitable for this task. Non-negative matrix factorization (NMF) produces non-negative sparse decompositions whose coefficients may be bimodally distributed but not binary as we require. We overcome this limitation by discretizing the matrix coefficients after factorization based on their distribution. Imposing additional explicit sparsity constraints on the NMF projection, e.g., using the approach of Gao et al. [22] did not improve the representation in any significant way.

The LEM approach we have described results in the generation of a small number of coregulated sub-signatures of genes. Several reference collections of coordinately expressed groups of genes are currently available for the analysis of gene expression data. For instance, an analysis of 246 subsets of mouse immune cells identified modules of genes with coordinated expression across diverse lineages, and was able to infer regulatory mechanisms controlling particular modules of genes [23]. In the human immune system, several studies have applied similar approaches to identify groups of co-regulated gene modules from expression profiles derived from PBMC samples representing a range of states of health and disease [24,25]. These PBMC modules have proven to be powerful tools for analyzing the human PBMC transcriptional response to infection, autoimmunity, and vaccination. However the difference with these existing collections is that the statistical interdependence of the genes in a particular module is defined a *priori* and the module collections are context dependent and static. These approaches assume that that interdependence of gene expression in the previously defined modules will be maintained in all future experiments, an assumption that has not been exhaustively tested. In contrast, our approach adaptively defines the association of genes based on their co-regulation in the experimental dataset being studied. Context-specific differences in the co-regulation of genes will therefore be captured by our approach, allowing the construction of a set of LEMs tailored to the specific experimental setting of interest. Additional studies will help define the extent of variation in the structure of LEMs from one biological context to another.

We applied our approach to defining LEMs to the problem of identifying features of the transcriptional response to vaccination. We showed that different vaccines elicit distinct kinetics of gene expression changes at day 3 and day 7 post vaccination compared to the prevaccination state. Vaccines such as YFV show marked similarity in gene expression at both time points, while MCV elicits a pattern of LEMs at day 3 that is quite distinct from that seen at day 7. This difference in the progression of biological changes elicited by vaccines underscores the difference in the biologic basis by which protein-conjugate and live viral vaccines elicit protective immune response. Consistent with this, the profile of LEMs elicited at day 3 by the five vaccines studied showed clear differences between vaccines comprising of protein-conjugates (MPSV and MCV4) and those derived from viruses. However, by day 7 the pattern of LEMs elicited by the different viruses started to converge. This suggests that while different mechanisms may be responsible for the initial events in the priming of an immune response, common patterns of immune response begin to emerge at later time points. These findings have implications for the point at which gene-expression-based predictors of vaccine response should be measured.

## Methods

### Overview

The goal of the LEM algorithm is to identify component biological sub-signatures present in a group of gene sets that are enriched in a phenotypic comparison of interest. This algorithm starts with *m* gene sets from a GSEA analysis and yields a small number of groups of genes, which we term leading edge metagenes (LEMs), that capture the biological processes differentially present in the phenotypic comparison. There are five key elements in this method:

#### Step 1: Identify enriched gene sets and their leading edges

We first identify gene sets that are enriched in a phenotypic comparison of interest, such as gene expression profiles of PBMC samples before and after vaccination. GSEA has been extensively described [6] and typically queries a list of genes ranked by their differential expression in two phenotypes with gene sets from databases such as MSigDB [9]. GSEA calculates an enrichment score (ES) that reflects the degree to which a set of genes is overrepresented at the extremes (top or bottom) of the ranked list. The statistical significance of this overrepresentation is estimated using an empirical phenotype-based permutation test. Enriched gene sets are considered to be those that exceed a statistical threshold set by the user, and can, in typical experiments include dozens or even hundreds of gene sets. The leading edge subset of genes in an enriched gene set are defined as those which appear in the ranked list before the point at which the running sum of the ES is greatest ([6], [Fig. S3). Leading edge genes therefore represent the most enriched subset of genes in a gene set.

#### Step 2: Construct the leading edge sparse matrix

We consolidate the leading edges of the *m* top-scoring gene sets into a sparse *n* by *m* matrix *M*, where the number of rows is the cardinality of the union of genes from all the leading edges in the *m* top gene sets, and the columns correspond to the genes in the m enriched gene sets. The value of each entry in the matrix is the signal to noise ratio of the corresponding gene between two conditions in comparison (Eq. 1) and 0 if the gene is not in the leading edge of that gene set. A large signal to noise ratio indicates a great difference in the gene’s expression between the two conditions.

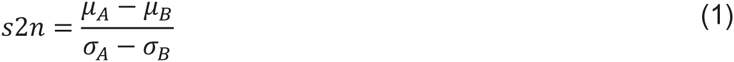

#### Step 3: Estimate the number of clusters in the leading edge sparse matrix via NMF

We use nonnegative matrix factorization (NMF) to identify clusters of genes with a similar pattern of leading edge membership in multiple gene sets and up‐ or down-regulation in the phenotype of interest. NMF is an efficient method for identifying hidden structure within a dataset [26]. Here we use NMF coupled with a model selection mechanism to estimate the number of clusters within the leading edge sparse matrix. Specifically, for a given leading edge sparse matrix *M* with *n* rows and *m* columns, we can approximate the original gene sets as positive linear combinations of these metagenes. The described procedure is equivalent to factoring the matrix *M* into two matrices such that *M* ≈ W × H [10,26]. The *W* matrix is a low-dimensional (rank *k* and *k* ≪ *m*) representation of the *M* matrix and each dimension of *W* is a positive linear combination of *n* genes, termed a metagene. Each entry in the *W* matrix represents the contribution of each gene to the metagene and each entry in the *H* matrix represents the amount of each metagene required to recapitulate the gene’s expression in each of the m gene sets. To find the best rank *k* of the *W* matrix, we use a consensus clustering method framework as previously described [11,27].

#### Step 4: Adaptively filter coefficients in the *W* matrix

Inspection of the *W* matrix shows that in each metagene, the coefficients of most genes are very small, and only a small number of genes have a coefficient significantly larger than 0. As each metagene is a positive linear combination of all the genes, a small coefficient indicates negligible contribution to the metagene. Thus the next step of our algorithm involves filtering out genes with small coefficients in each metagene. To do this, we first assume that the background distribution of coefficients fulfills an exponential distribution (S2 Fig.). We set a filtering threshold at the 95% quantile of the fitted exponential distribution and set all coefficients below this threshold to zero.

#### Step 5: Assign genes to the leading edge metagenes

As each gene can contribute to more than one metagene, we next need to assign each gene to a single metagene. The assignment of genes to metagenes uses the following rules: 1) if one gene has no contribution to any of the metagenes, it is not included in any metagene; 2) each gene with a coefficient above the threshold defined in Step 4 is assigned to the metagene in which it has the largest coefficient.

### Data preprocessing

We analyzed 5 existing datasets of gene expression profiles of PBMC from vaccinated subjects at three time points, day 0, day 3 and day 7 respectively. The GenePattern module “CollapseDataset” was used to extract the expression values of genes from the raw data file and to map Affymetrix probes to gene symbols [28]. We then applied quantile normalization and a log2 transformation to the expression data. The GEO accession ID of the five datasets are: GSE52245 for MPSV4 and MCV4; GSE13485 for YF-17D; GSE29617 for TIV and GSE29615 for LAIV [12,13,14].

### Gene set enrichment analysis

GSEA yields an enrichment score (ES) to quantify the overrepresentation of a set of genes *S* (e.g. genes coordinately up‐ or down-regulated in previous experiments) at the top or bottom of a ranked list of genes *L*. Candidate genes are ranked by their differential expression between two phenotypes, day 0 vs. day 3 and day 0 vs. day 7 in our case. The statistic is a weighted Kolmogorov-Smirnov-like statistic and significance is calculated using an empirical permutation test [6]. We used the desktop version of the GSEA software (http://www.gsea-msigdb.org) to conduct the leading edge analysis and extract the leading edge sparse matrix [6]. The GSEA software recommends an FDR < 0.25 as the cutoff to select significantly enriched gene sets for leading edge analysis. The number of gene sets satisfying this criterion ranged from 0 to 600 depending on the data set. The number of gene sets satisfying this criterion was less than 5 for six of the GSEA analyses, 186 for another, and between 186 and 550 for the remaining three. Thus, to maintain consistency in metagene detection, we used 500 as the number of top scoring gene sets for leading edge analysis for all the GSEA results.

### Calculate frequency of genes in the leading edges

After generating a leading edge sparse matrix, we enumerated the frequency of each gene in all leading edges by converting the leading edge sparse matrix into a binary matrix and summed up the values in each column.

### Annotate biological themes of the leading edge metagenes

To systematically annotate all the metagenes generated in our analysis, we used the DAVID annotation tool together with REVIGO [5,17,29] to calculate the overrepresentation of genes related to specific biological processes in each of the metagenes.

### Circos plots

To visualize the overlap in gene membership between metagenes, we generated Circos plots based on the number of genes in each metagene and the number of genes shared by two metagenes (http://mkweb.bcgsc.ca/tableviewer/).

### Calculate the significance of overlaps between metagenes

We used Fisher’s exact test to assess the significance of the number of genes shared by two metagenes. For example, to test the significance in gene sharing between Metagene 1 of LAIV at day 3 (M1.D3) versus Metagene 1 of LAIV at day 7 (M1.D7), we constructed a contingency table. Under the null hypothesis, the probability of obtaining a shown set of values follows a hypergeometric distribution:

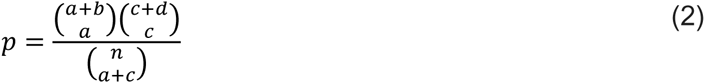

Where n = a + b + c + d

|  | Genes in M1.D3 | Genes not in M1.D3 |
| --- | --- | --- |
| Genes in M1.D7 | a | b |
| Genes not in M1.D7 | c | d |

### Calculate the distance between two gene signatures based on metagenes

Each phenotypic comparison (e.g., gene expression profiles of PBMC at day 0 versus day 3 post-vaccination) generates a set of metagenes that represents a unique signature associated with that comparison. To calculate the distance between two gene signatures, we first compute the *P*-values of Fisher’s exact test for each pairwise comparison of metagenes. The distance is then calculated as the sum of all the *P*-values normalized to the total number of pairs [30]. We used this metric as an input for hierarchical clustering.

## Data availability

The Leading Edge Metagene analysis method and source code are available as a Git repository hosted on GitHub (https://github.com/lamarck2008/LEM)

## Supplementary Figure Legends

**Supplementary Figure 1.**
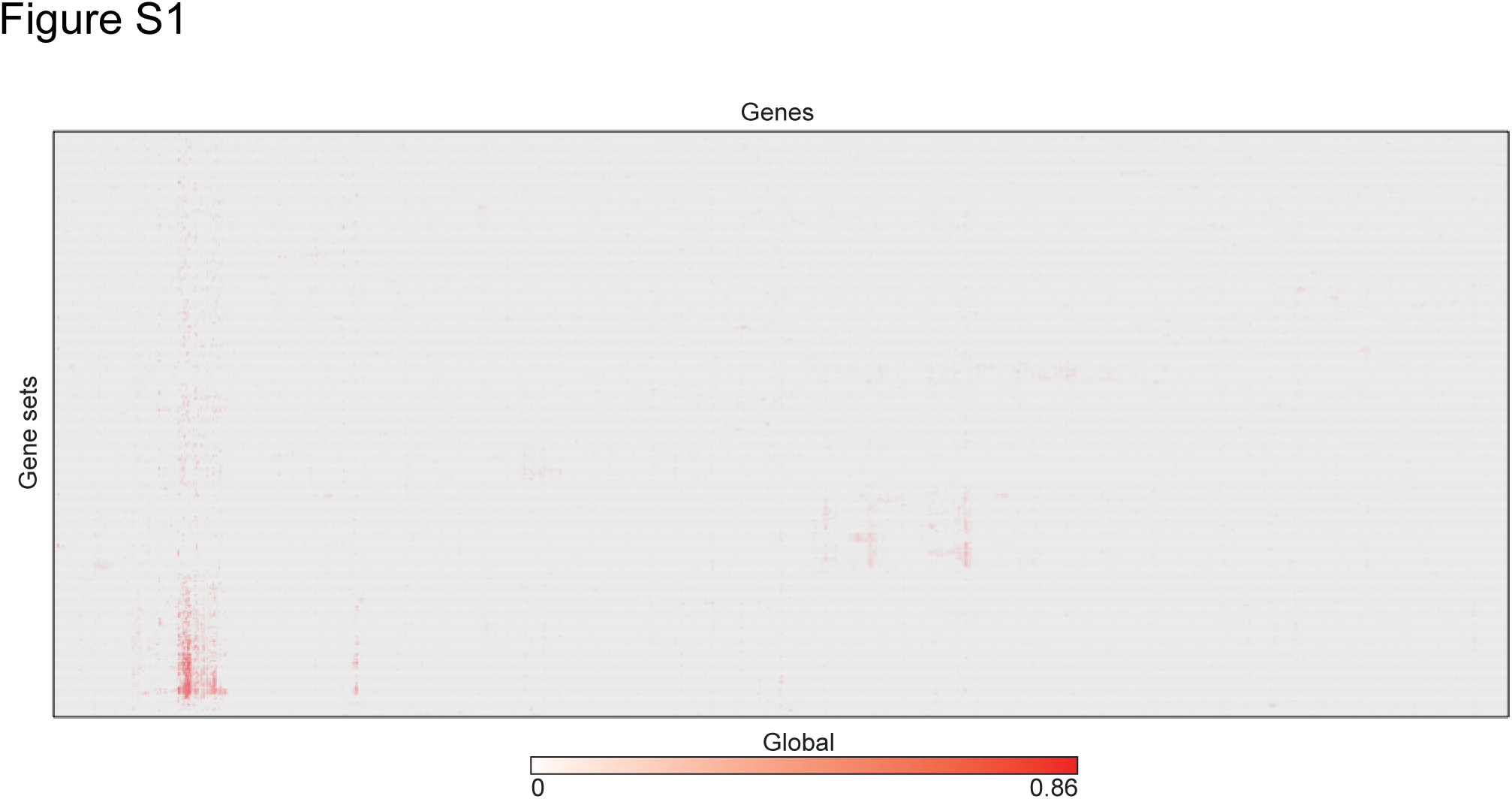
Leading edge sparse matrix. The genes by gene sets matrix generated by taking the leading edge analysis results of gene sets significantly enriched in PBMC samples of subjects 7 days after vaccination with YF-17D.

**Supplementary Figure 2.**
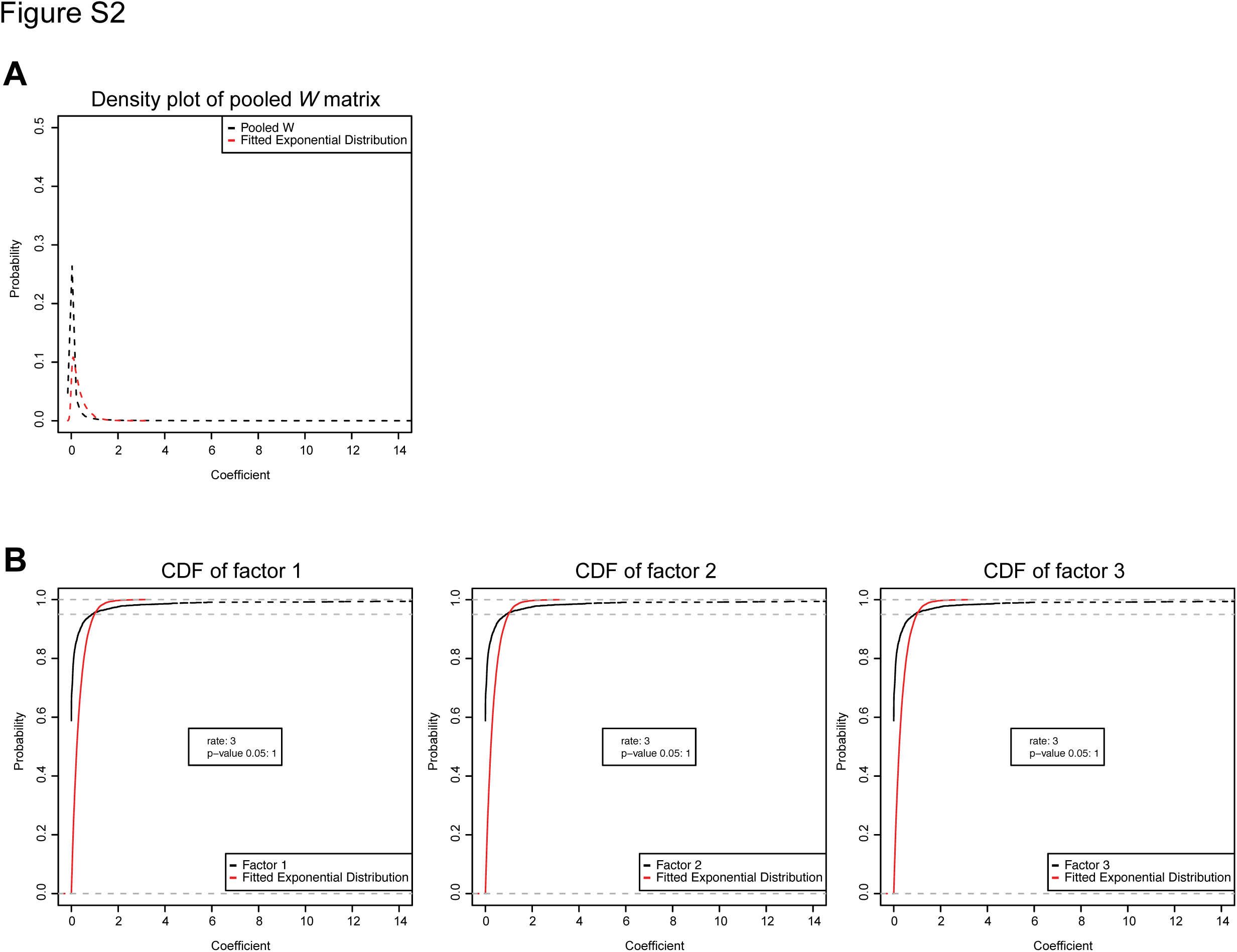
Distribution of coefficients in *W* matrix and fitted exponential distribution. **(A)** Density distribution of coefficients in *W* matrix (black dashed line) and fitted exponential distribution (red dashed line). **(B)** Accumulative distribution of coefficients in each column of *W* matrix (black dashed line) and the corresponding fitted exponential distribution (red dashed line).

**Supplementary Figure 3.**
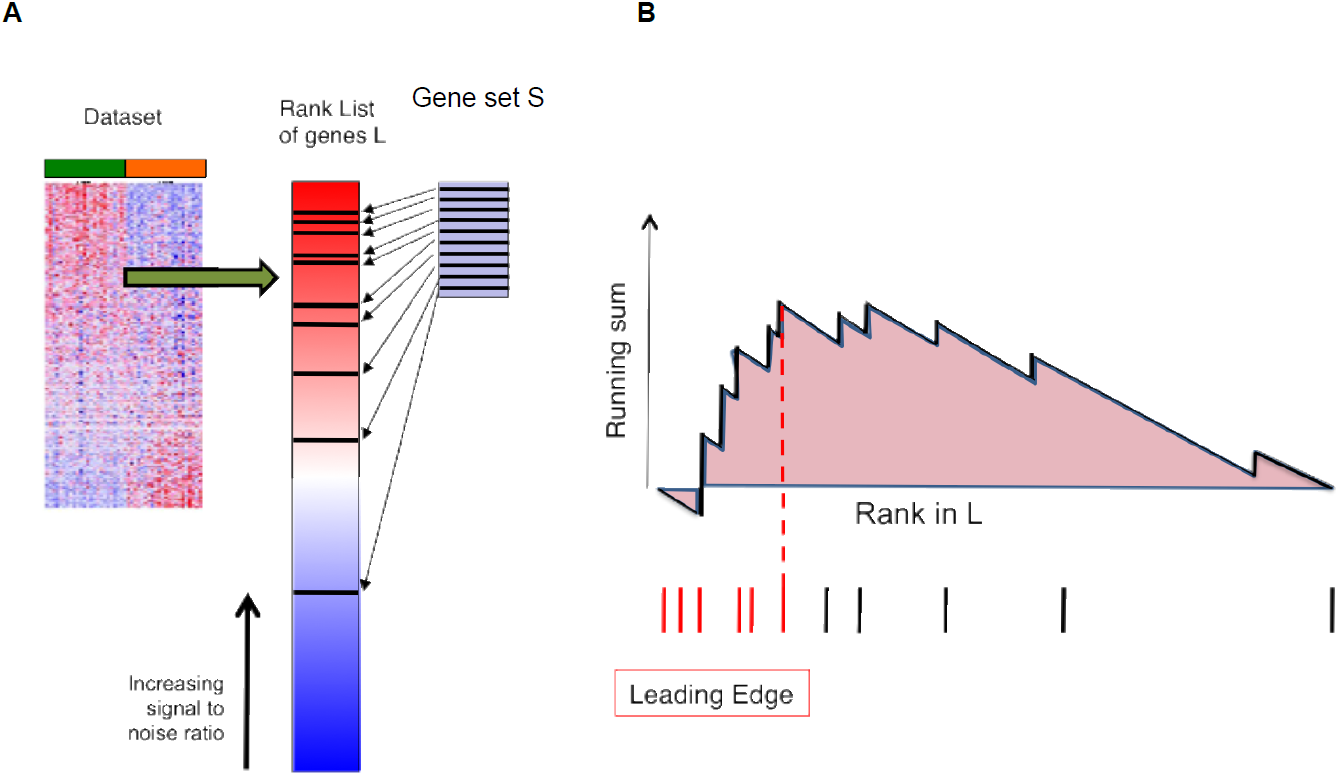
Gene set enrichment analysis schema and Leading edge subset of gene sets. **(A)** GSEA yields a quantitative measure of the overrepresentation of a set of genes *S* (e.g. genes encoding products in a same metabolic pathway) at the top or bottom of a ranked list of genes *L*. Genes in *L* is ranked by their signal to noise ratio with respect to the phenotype of comparison. **(B)** A running sum is calculated when computing the enrichment of each gene set. Leading edge subset of a gene set is highlighted in red. It corresponds to those genes in the gene set that appear in the ranked list *L* at, or before, the point where the running sum reaches its maximum deviation from zero, i.e., the Enrichment Score for that gene set.

